# A Novel 3D Imaging Approach for Quantification of GLUT4 Levels across the Intact Myocardium

**DOI:** 10.1101/2024.03.26.586764

**Authors:** Angéline Geiser, Susan Currie, Hadi Al-Hasani, Alexandra Chadt, Gail McConnell, Gwyn W. Gould

## Abstract

Cellular heterogeneity is a well-accepted feature of tissues, and both transcriptional and metabolic diversity have been revealed by numerous approaches, including optical imaging. However, the high magnification objective lenses needed for high-resolution imaging provides information from only small layers of tissue, which can result in poor cell statistics. There is therefore an unmet need for an imaging modality that can provide detailed molecular and cellular insight within intact tissue samples in 3D. Using GFP-tagged GLUT4 as proof of concept, we present here a novel optical mesoscopy approach that allows precise measurement of the spatial location of GLUT4 within specific anatomical structures across the myocardium in ultrathick sections (5 mm x 5 mm x 3 mm) of intact mouse heart. We reveal distinct GLUT4 distribution patterns across cardiac walls and highlight specific changes in GLUT4 expression levels in response to high fat diet-feeding, and we identify gender-dependent differences in expression patterns. This method is applicable to any target that can be labelled for light microscopy, and to other complex tissues when organ structure needs to be considered simultaneously with cellular detail.

**SUMMARY STATEMENT:** Here we present a novel 3D optical mesoscopy approach that allows the study of both GLUT4 protein expression levels and structural distribution within ultrathick sections of intact murine hearts, in response to high fat diet-feeding.

## INTRODUCTION

The unique energy-demand of the heart means that there is a complete turnover of myocardial ATP every 10 seconds at rest. To accommodate this, the heart can generate energy from lipids, carbohydrates, lactate, amino acids, as well as ketone bodies, depending on the prevailing metabolic milieu. It is this ability to adapt and shift between different substrates, ensuring a continuous energy supply, that has earned the heart the name of “metabolic omnivore” (Taegtmeyer et al., 2004). This flexibility plays a key role in physiological cardiac function, enabling the heart to adapt to specific energy-demands, substrate availability, as well as stress conditions such as increased pressure load, injury, or ischemia (Stanley et al., 2005).

Over the years, the identification of metabolic abnormalities in heart failure (HF) has driven numerous studies to better define metabolic differences between healthy and diseased hearts. The use of *in vivo* imaging approaches has provided a wealth of insight into cardiac metabolic activity, for example, by quantifying absolute values of ATP and phosphocreatine (PCr) using ^31^P magnetic resonance spectroscopy. Studies have reported reduced PCr:ATP ratios in patients with HF, confirming metabolic dysfunction (Conway et al., 1991; Hardy et al., 1991; Neubauer et al., 1992; Neubauer et al., 1997; Tsampasian et al., 2023). Similarly, positron emission tomography (PET) was developed as a unique non-invasive approach to evaluate myocardial glucose and fatty acid turnover, as well as tissue perfusion (Taylor et al., 2001; Kudo et al., 2002). PET can quantify regional myocardial blood flow, providing functional and structural insights on the effects of different diseases across distinct anatomical regions of the heart, as well as revealing changes in both global and regional cardiac metabolism. However, the spatial resolution of these methods is poor, and the mechanisms behind these metabolic changes remain elusive at the single-cell level and challenging to quantify in individual cells within the intact heart.

Differences between atrial and ventricular metabolism have been established in numerous mammalian species (Bass et al., 1993). In addition, considering the major risk factors of HF, which includes diabetes and obesity (Kannel et al., 1974; Kenchaiah et al., 2002; Kenny and Abel, 2019), studies have described distinct heterogenous features associated with such diseases (Edvardsen and Klaeboe, 2019). The hemodynamic changes observed in obesity have been shown to induce left atrial enlargement as well as left ventricular wall stress (Alpert et al., 2014). Similarly, the altered metabolic milieu seen in diabetes has been shown to contribute to cardiac chamber remodelling. For instance, Gulsin et al. (2019) showed that patients with HF with preserved ejection fraction and type 2 diabetes (T2D), have increased concentric left ventricular remodelling, decreased left atrial volumes, and increased systemic inflammation, when compared to patients without T2D. Linssen et al. (2020) also associate (pre)diabetes with structural changes in the right atria and right ventricle, as well as impaired right ventricular systolic and diastolic function. Cardiac remodelling is a multifactorial process that not only includes a structural response of myocardial tissue to different pathophysiological processes (Cohn, 1995; Planinc et al., 2021), but also involves biochemical remodelling (van Bilsen et al., 2004). However, linking the two, and more specifically, linking them to distinct metabolic changes is challenging, as there remains an unmet need to quantify protein levels and changes within distinct three-dimensional (3D) strucures of intact hearts. This is further emphasised by *in vitro* studies such as those performed by Doll et al. and Linscheid et al., who produced quantitative proteomic maps of human hearts (Doll et al., 2017; Linscheid et al., 2020). These studies provide cardiac metabolic information at a molecular level, highlighting regional variations in protein expression within different heart regions, and that this may be reorganised in diseased hearts.

Focusing on the major glucose transporter expressed in the heart, GLUT4, Ware et al. (2011) observed a 2.5-fold increase in protein content in the left ventricle in chronic HF. This up-regulation was suggested to provide an adaptive metabolic profile to support increased energetic demands in response to ventricular wall stress and hypertrophy. They also detected a paradoxical decrease in GLUT4 content in the right atria (Ware et al., 2011). Similarly, Maria et al. (2015) observed a down-regulation of GLUT4 levels by 70% at the surface of cardiomyocytes in T1D atria which they postulated might limit the ability of the heart to properly use glucose as a source of energy in diabetes, therefore impairing recovery (Maria et al., 2015). In addition, distinct patterns of GLUT4 and GLUT8 expression in different heart chambers highlight a further aspect of cardiac regional heterogeneity, indistinguishable in clinical studies, whereby different GLUT isoforms may play a greater role in regulating glucose uptake in the myocardium under pathological circumstances (Maria et al., 2015). However, it is unclear whether these reported variations in specific protein expression represent a global change, or whether distinct architectural features within different regions of the heart adapt differently to pathological stimuli. To date, most approaches for the study of regional protein heterogeneity rely on some degree of manual tissue dissection, thus introducing a level of error, inconsistency, and destruction of specific regional information which could be evident in intact tissues. Such data underscores the need for detailed studies focussing on the expression of specific proteins within the 3D structure of the intact heart, prompting us to consider how *organ-scale* analysis of protein expression patterns could be developed.

Here we present a novel optical mesoscopy approach that allows the study of both GLUT4 protein expression levels and structural distribution within ultrathick sections of intact murine hearts. As the primary glucose transporter expressed in the heart, GLUT4 represents a key component of glucose homeostasis (Katz et al., 1995; Abel, 2004; Szablewski, 2017; Wende et al., 2017), hence we use in this study GFP-tagged GLUT4 as proof of concept. We show how regional variations in GLUT4 expression can be observed and quantified systematically in whole mounts of heart tissue up to 5 mm x 5 mm x 3 mm in size, while allowing a clear view of intact protein levels within this large tissue volume. We also compare GLUT4 distribution in intact hearts from mice fed standard chow or high-fat diet, the latter being a well-established method to induce obesity and insulin resistance (Inui, 2003; Speakman et al., 2007; Sato et al., 2010). We use this model to ascertain how GLUT4 levels and distribution may be affected under pathological circumstances caused by high-fat diet. The methodology and analysis pipelines developed here clearly reveal the capacity of novel molecular insight on the mesoscale using this approach. This method may be useful for the analysis of other markers in studies of cardiac disease and can be readily adapted to other complex tissues when organ structure needs to be considered with high spatial resolution.

## RESULTS

Here we used transgenic mice expressing HA- and GFP-tagged GLUT4 (Fazakerley et al., 2009; Lizunov et al., 2012). This transporter construct is widely used in studies of GLUT4 trafficking and is known to exhibit insulin-dependent movement to the plasma membrane in response to insulin in isolated cells, including cardiomyocytes (Blot and McGraw, 2008; Muretta et al., 2008; Fazakerley et al., 2009; Habtemichael et al., 2011; Klip et al., 2019). Hence, we reasoned that this would provide a useful proof of concept for our analysis pipeline, avoiding immunostaining in the development of this approach.

In this study, we use the term intact to reflect the use of isolated heart sections, typically 5 mm x 5 mm x 3 mm in size, where the core structure remains untouched and individual cardiac chambers undissected (for details see Methods).

### Axial Imaging Depth with and without Organ Perfusion

Using optical mesoscopy, we were able to visualise GFP-tagged GLUT4 within optically cleared ultrathick sections of intact murine hearts (Fig. 1A). Samples were cleared using the iDISCO method (Renier et al., 2014) (for details see Methods). Fig. 1B displays an average intensity axial-projection of GLUT4-GFP fluorescence within a whole mount (5 mm × 5 mm x 3 mm) of a typical ventricular cardiac tissue section. Visible intact 3D anatomical structures including the left ventricle (LV), the two papillary muscles of the LV, the interventricular septum, as well as the right ventricle (RV) are clearly observed (Fig. 1B).

**Figure 1.**
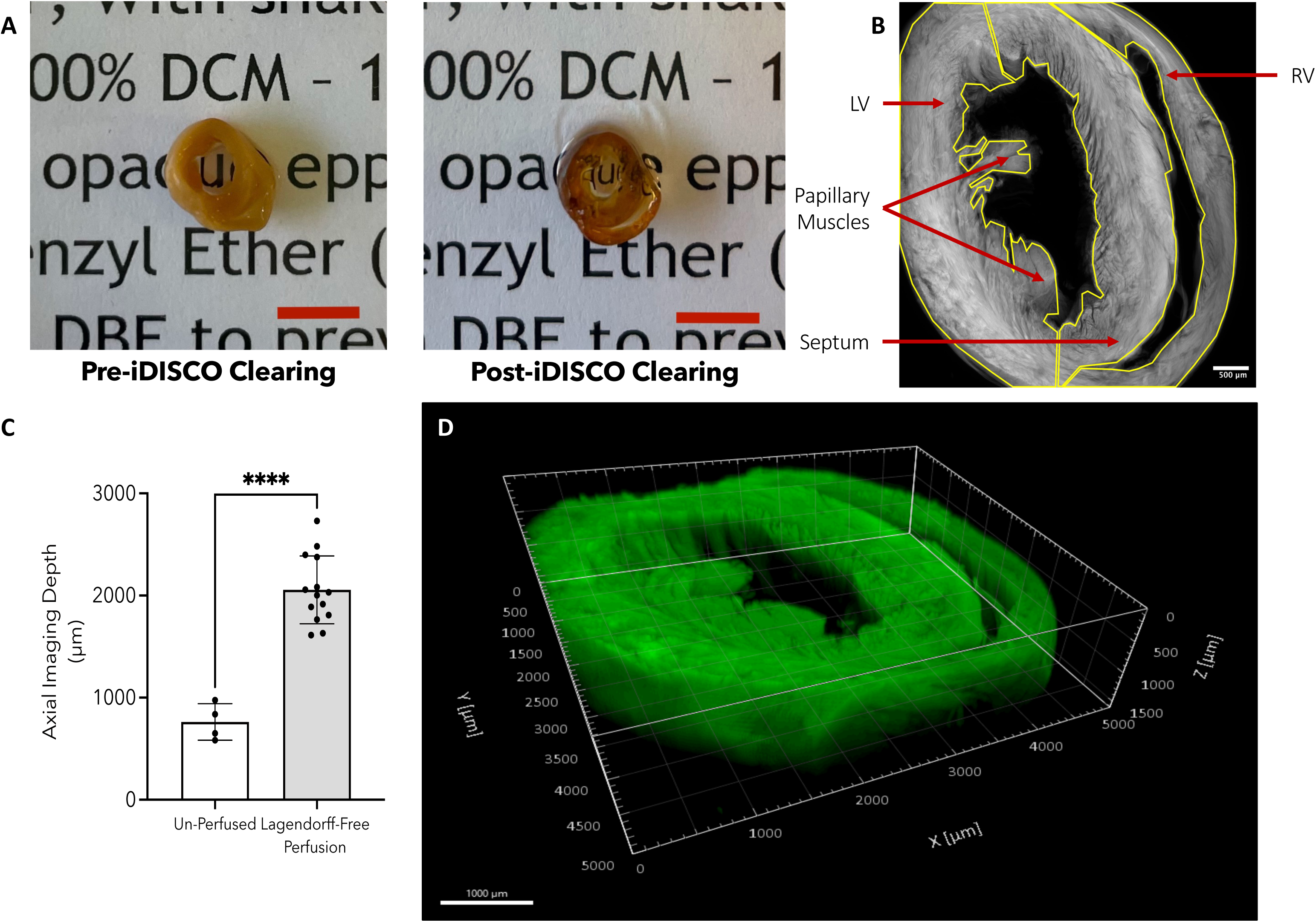
Visualisation of GLUT4-GFP in ultrathick sections of cleared mouse heart and characterisation of axial imaging depth with and without organ perfusion. (**A**) Representative images of pre- and post-iDISCO optical clearing of a perfused 3-mm thick mouse ventricular heart section. Scale bar (red) = 5 mm. (**B**) Representative average intensity axial-projected image of GLUT4-GFP fluorescence within perfused and iDISCO optically cleared 3-mm thick mouse ventricular heart section showing distinct structural features: the LV, the two papillary muscles of the LV, the interventricular septum, and the RV. Scale bars = 500 μm. (**C**) Mean ± s.d. of axial imaging depth (µm) in un-perfused (white-bar; n = 4) iDISCO optically cleared 3-mm thick mouse ventricular heart sections compared to perfused (grey-bar; Langendorff-Free Perfusion, n = 14) hearts. * indicates a significant increase in axial imaging depth upon Langendorff-Free perfusion (2.06 ± 0.33 mm) compared to un-perfused (0.76 ± 0.18 mm) hearts, p ≤ 0.05 (Unpaired t-test with Welch’s correction). (**D**) 3D render of GLUT4-GFP fluorescence (green) within a whole mount of optically cleared 3-mm thick mouse ventricular heart section. Scale bars = 1000 μm. Specimens were imaged using the Mesolens system over a 5 mm × 5mm x 3mm volume using a z-step size of 5μm as described in Methods.

Optimisation of the tissue clearing protocol has highlighted the importance of blood removal for organ optical clearing, especially in heme-rich tissues such as the heart. Simple buffer perfusion of the tissue was shown to effectively remove unwanted red blood cells. As described in the Methods, a Langendorff-free perfusion method was used to remove as much blood as possible from isolated hearts. Without this perfusion step, a total imaging depth of heart sections was proven to be restricted to less than 1 mm (0.76 ± 0.18 mm; Fig. 1C). With the inclusion of the perfusion step in our method, specimens (n = 14 included in this study) were imaged using a 5 μm z-step size and yielded an average imaging depth of 2.06 ± 0.33 mm (Fig. 1C,D and Movie S1).

### Quantitative Analysis of GLUT4 Levels and Distribution within Intact Ventricular Cardiac Sections

Molecular and structural regional heterogeneity, in both healthy and diseased hearts, have become increasingly characterised (Bass et al., 1993; Alpert et al., 2014; Maria et al., 2015; Doll et al., 2017; Gulsin et al., 2019; Linscheid et al., 2020; Linssen et al., 2020). However, linking specific metabolic changes and protein expression patterns to the 3D architecture of intact heart regions remains an unmet challenge. Considering that the fluorescence of GLUT4-GFP within ultrathick cardiac tissue section can be effectively detected using optical mesoscopy (Fig. 1), we therefore reasoned that regional variations in GLUT4 expression could be observed and systematically quantified within acquired 3D reconstructed volumes of intact ventricular heart sections. We chose to test this hypothesis using hearts from mice fed standard chow (Control) or high-fat (HFD) diets to examine how pathophysiological changes may be identified using our method. As expected, mice fed a HFD presented with elevated body weight and blood glucose (Fig. S2A,B,C) (Inui, 2003; Speakman et al., 2007; Sato et al., 2010).

The developed analysis pipeline of image processing, visualisation, and quantification of GLUT4-GFP fluorescent signals is outlined in detail in the Methods and summarised in Fig. 2.

**Figure 2.**
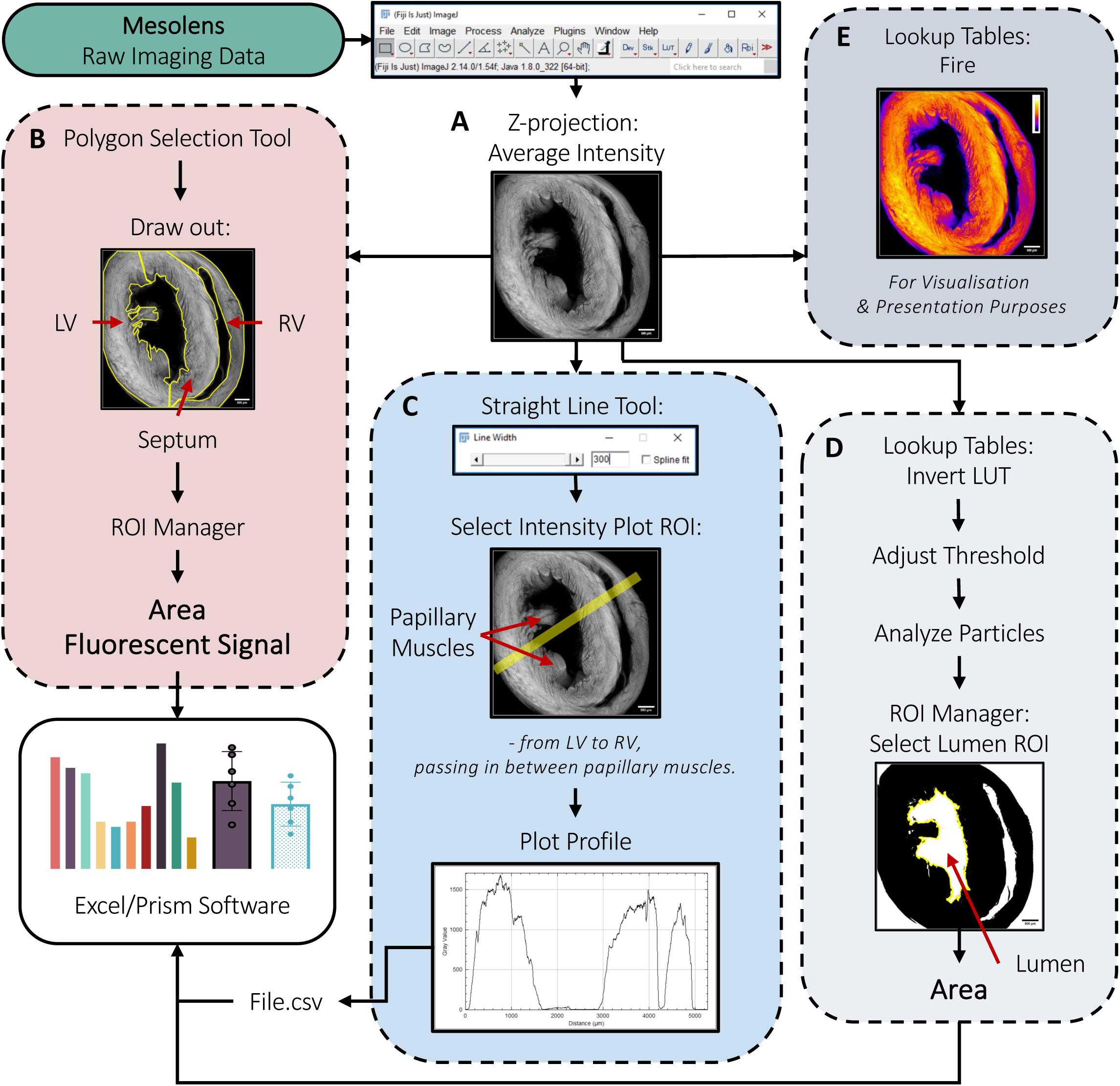
Summarised flowchart of analysis pipeline for quantification of GLUT4-GFP levels in ultrathick sections of cleared mouse heart. Following acquisition, raw Mesolens imaging data were processed using ImageJ image processing software (Schindelin et al., 2012). (**A**) Acquired 3D volumes were transformed as 2D images by performing an average intensity axial-projection. (**B**) Using the polygon selection tool, ROIs within heart sections, including the LV, septum, and RV, were manually segmented, and saved in the ROI Manager to then allow measurement of the surface area and GLUT4-GFP fluorescent signals. (**C**) Using the line selection tool, intensity plot ROI was manually segmented by drawing a line from the LV to the RV, crossing the septum and passing in between LV papillary muscles. The Plot Profile tool was then used to generate fluorescence intensity profiles. (**D**) To isolate and quantify the area of the lumen, axial-projected datasets were subjected to thresholding. The Analyze Particles tool was then used to the lumen ROI. Area of the lumen was then measured. (**E**) Colour-coding of fluorescence intensity within axial-project images for presentation and visualisation purposes was performed using the “fire” lookup table. Extracted values and parameters from ImageJ were further processed as described in the Methods, together with the procedures for raw Mesolens imaging data processing, visualisation, and quantification.

### Global Regional GLUT4 Expression Levels

To quantify and compare regional GLUT4-GFP protein content in both Control and HFD intact ventricular heart sections, we first displayed the Mesolens-acquired reconstructed volumes as 2D images by performing an average intensity axial-projection of the 3D datasets. All subsequent analysis were then performed from each specimen axial-projected images (Fig. 2 and see Methods for more details). Fig. 3A displays representative axial-projections of GLUT4-GFP fluorescence within Control and HFD hearts. In both groups, no statistically significant differences in GLUT4-GFP fluorescence, expressed relative to left ventricular signals, were observed between the LV, septum, and RV (Fig. 3B). This result is in good agreement with immunoblotting data shown in Fig. 3C,D, where no significant regional heterogeneity in GLUT4-GFP levels is observed in tissue homogenates dissected from hearts of mice fed a standard chow diet. Nevertheless, this approach allows an appreciation of intrinsic ventricular and atrial regional heterogeneity (Fig. 3D) in protein expression evident within a healthy heart (Bass et al., 1993; Doll et al., 2017), facilitating a comparison of intact tissue in two distinct physiological states.

**Figure 3.**
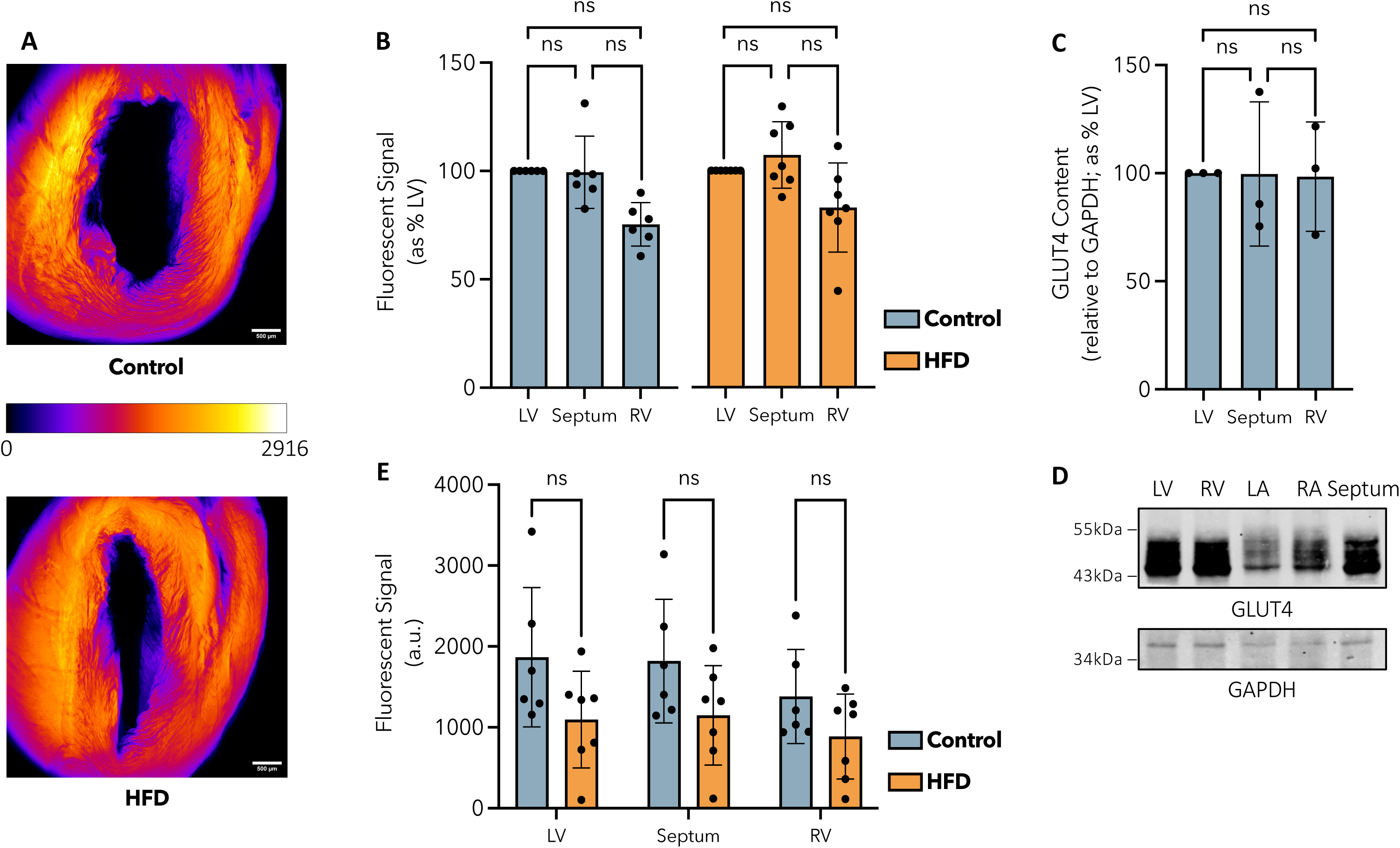
Representative axial-projections of GLUT4-GFP fluorescence and quantification of global regional GLUT4 expression levels in Control and HFD hearts. (**A**) Representative average intensity axial-projections of GLUT4-GFP fluorescence within Control and HFD hearts (Hearts No.5 and 11, respectively; Fig. S2D). Specimens were imaged over a 5 mm × 5mm x 3mm volume using a z-step size of 5μm as described in Methods. Axial-projected images were colour-coded by fluorescent intensity using the “fire” lookup table. Intensity values are expressed relative to the highest intensity values between the two images (maximum intensity = 2916.00 a.u.; Hearts No.5, Control). Scale bars = 500 μm. (**B**) Mean ± s.d. of GLUT4-GFP fluorescent signal in the LV, septum, and RV of Control (blue; n = 6) and HFD (orange; n = 7) hearts, expressed relative to % LV. Data indicate no statistically significant (ns) differences between cardiac regions within each group. Brown-Forsythe and Welch ANOVA tests were performed on raw data within each group. (**C**) Mean ± s.d. of total GLUT4 content in isolated LV, septum, and RV tissue homogenates from mice fed a standard chow diet. Values were quantified from immunoblots of the type shown in panel D, expressed relative to GAPDH, and compared to % LV (n = 3). Data indicate no statistically significant (ns) differences between cardiac regions (Brown-Forsythe and Welch ANOVA). (**D**) Representative immunoblot of GLUT4 levels in isolated heart sections from mice fed a standard chow diet, including the LV, RV, left atrium (LA), right atrium (RA), and the septum. GAPDH was used as a loading control; bands were obtained from the same membrane. Tissue homogenates were produced, analysed, and visualised as described in the Methods; data from a representative experiment are shown. (**E**) Mean ± s.d. of GLUT4-GFP fluorescent signal (a.u.) in the LV, septum, and RV of Control (blue; n = 6) hearts compared to HFD (orange; n = 7) hearts. Data indicate no statistically significant (ns) differences within each cardiac region between Control and HFD hearts (Multiple unpaired t-tests with Welch’s correction).

In Fig. 3E, we compare GLUT4-GFP signals within each studied cardiac region between Control and HFD hearts. While the data suggest a trend toward lowered GLUT4-GFP levels in HFD tissues, these differences did not reach statistical significance (Fig. 3E). We note that decreased GLUT4 in HFD-fed mice has been reported by others, highlighting the repressive effect of a HFD on total GLUT4 content in the heart (Wright et al., 2009; Jackson et al., 2015). To this end, it is also important to note that the GLUT4-GFP transgene in these animals is under the control of the muscle creatinine kinase promoter rather than the endogenous GLUT4 promoter, hence transcriptional control pathways may not be completely mimicked in this model. Nevertheless, these results illustrate a simple quantitative analysis procedure that enables the study of both global and regional protein content within anatomically complex tissue volumes.

### Transmural Fluorescence: GLUT4 Distribution Across Cardiac Walls

To assess protein distribution across the myocardium walls, we next quantified and compared transmural fluorescence profiles of GLUT4-GFP across the width (x-axis) of left ventricular, septal, and right ventricular walls within Mesolens-acquired 3D reconstructed cardiac volumes (Fig. 4). For clarity in the following communicated results, we note that all transmural datasets are displayed and were analysed, from left to right, in the flowing order: LV, septum, and RV.

**Figure 4.**
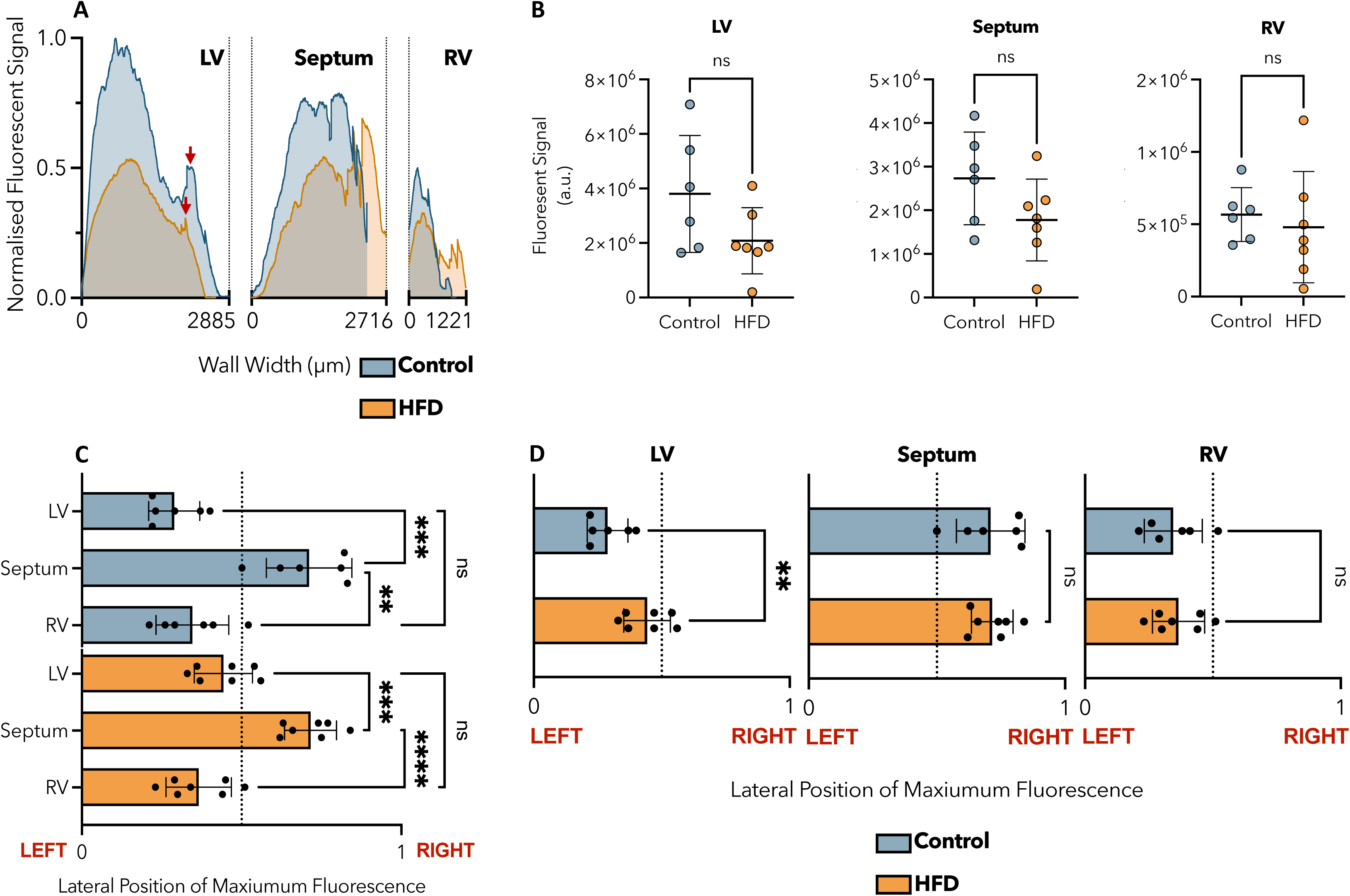
Quantification of transmural GLUT4-GFP fluorescence in Control and HFD hearts. (**A**) Mean transmural line profiles of GLUT4-GFP fluorescence across the width (x-axis) of left ventricular (LV), septal, and right ventricular (RV) walls in Control (blue; n = 6) and HFD (orange; n = 7) hearts. Fluorescent signal values are expressed relative to the highest intensity values between each group (maximum intensity = 2584.46 a.u.; Control, LV). Profiles were obtained as described in the Methods. The position of LV papillary muscles along line ROI profiles in both group is indicated by red arrows. (**B**) Mean ± s.d. of GLUT4-GFP fluorescent signal (a.u.) in the LV, septum, and RV of Control (blue; n = 6) hearts compared to HFD (orange; n = 7) hearts. Fluorescent signal values were measured from the area under curve of individual transmural fluorescence profiles as described in the Methods. Data indicate no statistically significant (ns) differences within each cardiac region between Control and HFD hearts. Unpaired t-tests with Welch’s correction were performed on raw data for each cardiac region. (**C**) Mean ± s.d. of GLUT4-GFP maximum fluorescence lateral position along the breadth of the LV, septum, and RV in Control (blue; n = 6) and HFD hearts (orange; n = 7), expressed relative to total wall width. In both Control and HFD hearts, data indicate GLUT4 fluorescence significantly peaking toward the right-hand side of the septal wall when compared to the LV and RV, where a left-skewed accumulation of GLUT4 proteins is observed along the width of both ventricular walls. **, ***, **** represent p ≤ 0.01, p ≤ 0.001, p ≤ 0.0001. Brown-Forsythe and Welch ANOVA tests were performed on raw data within each group. (**D**) Mean ± s.d. of GLUT4-GFP maximum fluorescence lateral position along the breadth of the LV, septum, and RV of Control (blue; n = 6) hearts compared to HFD (orange; n = 7) hearts, expressed relative to total wall width. Data indicate a significant change in the lateral position of GLUT4-GFP maximum fluorescence toward the centre of the LV in HFD hearts when compared to Control specimens. ** represents p ≤ 0.01. No statistically significant (ns) differences were observed between Control and HFD hearts within the septal and right ventricular walls. Unpaired t-tests with Welch’s correction were performed on raw data for each cardiac region. All lateral position values in C and D were measured from individual transmural fluorescence profiles as described in the Methods.

Cardiac regions were easily identified within extracted profiles (Fig.4A). Fig. 4A displays comparison of mean transmural fluorescence profiles of Control and HFD hearts. At first glance, the fluorescence intensity from GLUT4-GFP appears to be reduced in HFD hearts compared to Control specimens. However, similarly to previous results obtained from total regional measurements (Fig. 3E), quantification of fluorescent signals from the area under curve of each individual transmural profile revealed no statistically significant differences in any of the regions of study between Control and HFD mice (Fig. 4B).

In addition to total fluorescent signals, information on the position of where GLUT4-GFP fluorescence peaks along the breadth of each cardiac regions (lateral position of maximum fluorescence) were also extracted and normalised relative to total ventricular or septal wall width (Fig. 4C). Final values therefore illustrate whether detected GLUT4-GFP is expressed with a higher content toward the left or right-hand side of the studied ventricular or septal structures.

Our data highlight a significant regional heterogeneity in GLUT4-GFP transmural distribution amongst cardiac regions (Fig. 4C). In both Control and HFD hearts, Fig. 4C shows GLUT4-GFP fluorescence significantly peaking toward the right-hand side of the septal wall when compared to the LV and RV, where a left-skewed accumulation of GLUT4-GFP proteins is observed along the width of both ventricular walls (Fig. 4C). These results therefore suggest an asymmetry within GLUT4-GFP distribution profiles of different cardiac regions. Strikingly, while no differences were observed in the distribution of GLUT4-GFP within the septal and right ventricular walls, our data reveal a significant change in the lateral position of GLUT4-GFP maximum fluorescence toward the centre of the LV in HFD hearts when compared to Control specimens (Fig. 4D). Together, our data indicates that while total levels of GLUT4-GFP remain consistent in the LV, it appears that protein distribution across the left ventricular wall is changed upon HFD-feeding. This may well underscore changes in contractile activities in this region.

Finally, Fig. 4A also highlights the presence of GLUT4-GFP in more complex structural cardiac features: the LV papillary muscles (red arrow; Fig. 4A). Although not quantified in this analysis, this reveals the potential additional insight that our approach can provide and exemplifies the potential of using protein content distribution profiles across the width of the 3D structure under study.

### Gender-Specific Impact of a High-Fat Diet

No statistically significant differences were observed in GLUT4-GFP fluorescent signals and protein distribution patterns between male and female hearts, examined as a single cohort, within any of the examined cardiac sections (Fig. 5A,B). However, a significant decrease was observed in left ventricular GLUT4-GFP levels in the heart of HFD females compared to Control littermates (Fig. 5C). While the same statistical evaluation could not be applied to male specimens due to a low sample size for Control males (n = 2), these data suggest that a HFD over the course of 20 weeks may have a more significant impact on left ventricular GLUT4-GFP levels in females compared to males. No significant differences in GLUT4-GFP fluorescent signals between Control and HFD were found in the septum and RV of either males and females (Fig. S3G,H).

**Figure 5.**
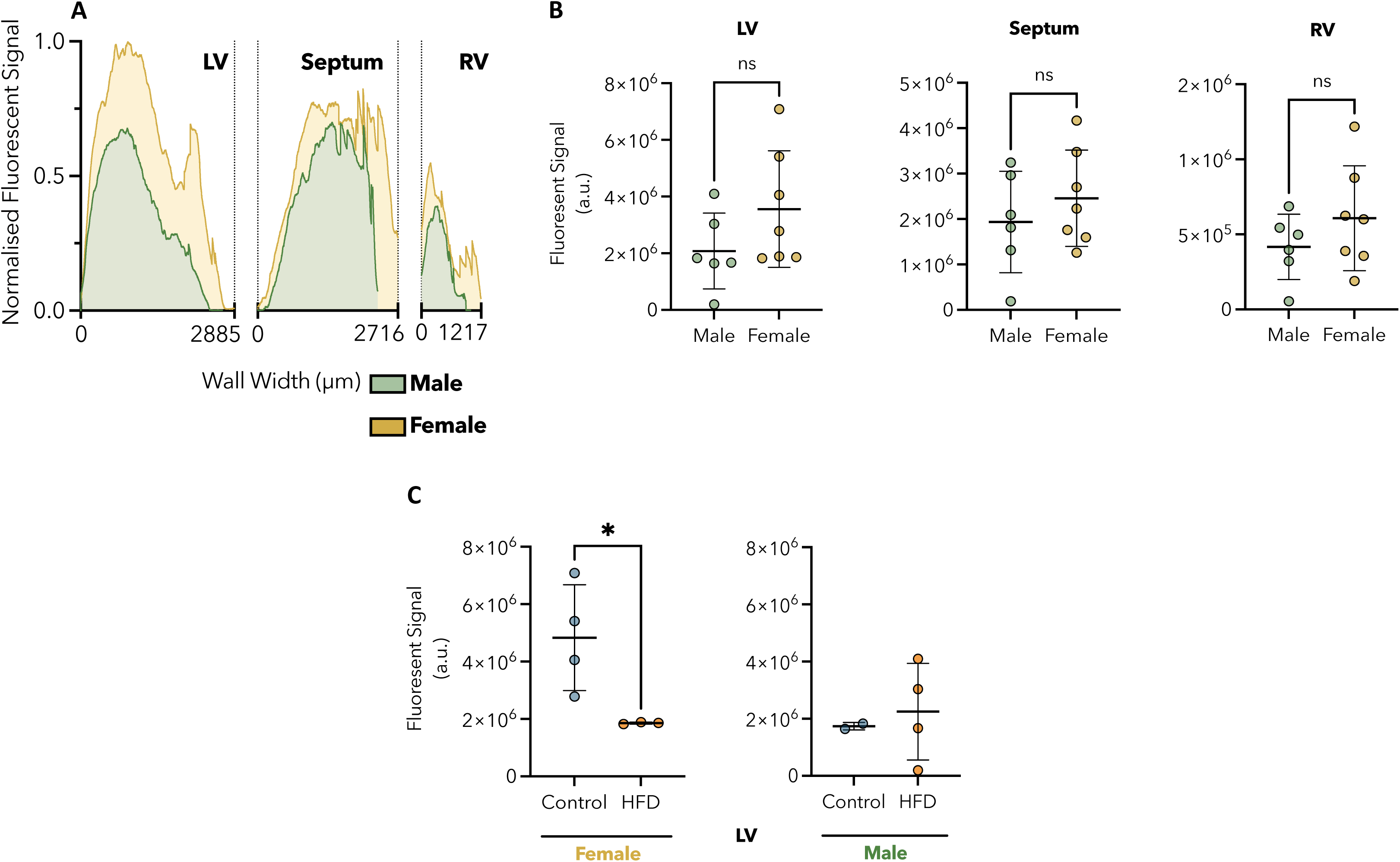
Quantification of transmural GLUT4-GFP fluorescence in male and female hearts and impact of HFD in females. (**A**) Mean transmural line profiles of GLUT4-GFP fluorescence across the width (x-axis) of left ventricular (LV), septal, and right ventricular (RV) walls in male (green; n = 6, Control = 2, HFD = 4) and female (yellow; n = 7, Control = 4, HFD = 3) hearts. Fluorescent signal values are expressed relative to the highest intensity values between each group (maximum intensity = 2236.20 a.u.; Female, LV). Profiles were obtained as described in the Methods. (**B**) Mean ± s.d. of GLUT4-GFP fluorescent signal (a.u.) in the LV, septum, and RV of male (green; n = 6, Control = 2, HFD = 4) hearts compared to female (yellow; n = 7, Control = 4, HFD = 3) hearts. Fluorescent signal values were measured from the area under curve of individual transmural fluorescence profiles as described in the Methods. Data indicate no statistically significant (ns) differences within each cardiac region between Control and HFD hearts. Unpaired t-tests with Welch’s correction were performed on raw data for each cardiac region. (**C**) Mean ± s.d. of GLUT4-GFP fluorescent signal (a.u.) in the LV of Control (blue) compared to HFD (orange) hearts, in female (yellow; n = 7, Control = 4, HFD = 3) and male (green; n = 6, Control = 2, HFD = 4). Fluorescent signal values were measured from the area under curve of individual transmural fluorescence profiles as described in the Methods. Data indicate a significant decrease in left ventricular GLUT4-GFP fluorescent signal in HFD female hearts when compared to Control littermates. * represents p ≤ 0.05 (Unpaired t-tests with Welch’s correction). No statistical evaluation was applied to male specimens due to a low sample size for Control males (n = 2).

In addition, a more variable and higher intra-specimen range in GLUT4 fluorescent signal is evident in Fig. 5C for Control females when compared to HFD littermates. Interestingly, the opposite can be observed in males, suggesting an inverted effect of HFD-feeding on left ventricular GLUT4 distribution between sexes. Further studies are needed to unravel potential mechanisms at play.

### Anatomical Parameters of Cardiac Structure

The 3D datasets can also be easily used to assess the gross anatomical features of the imaged myocardium. In addition to visualisation and measurement of fluorescent signals from GLUT4-GFP, we also extracted further anatomical parameters of cardiac structure from the Mesolens-acquired 3D reconstructed ventricular volumes, including the surface area and width of cardiac walls of each studied region, as well as the size of the lumen (Fig. S3A,B,C,D,E,F and Methods for details). While comparison of fluorescence between Control and HFD tissues showed no statistical significance, these results exemplify the ability of optical mesoscopy to assess protein levels in concert with quantifiable parameters of cardiac structures.

Overall, we provide here a set of quantitative analysis options allowing for better understanding of GLUT4 distribution within 3D specimens of intact hearts using the open-source image processing software ImageJ. This analysis pipeline remains widely adaptable to diverse molecular and structural target of interest.

## DISCUSSION

In this study, we used, for the first time, the imaging capability of the Mesolens to visualise and analyse intact cardiac tissues. We show that optical mesoscopy allows for a non-destructive visualisation of whole mounts of anatomical intact heart sections up to 5 mm x 5 mm x 3 mm in size with high spatial resolution in 3D. Using GFP-tagged GLUT4 as proof of concept, we demonstrate that the visualisation of GLUT4-GFP fluorescence within distinct anatomical structures across the myocardium, specifically the LV, the two papillary muscles of the LV, the interventricular septum, as well as the RV, is feasible in tissues up to 3-mm thick (Fig. 1). The use of a Langendorff-based perfusion method for the isolation of heme-rich organs such as the heart, has been suggested to improve axial imaging depth (Bell et al., 2011): the current limitation of imaging depth is the 3 mm working distance of the Mesolens (McConnell et al., 2016) but this presents an opportunity for further bioimaging technology development.

The presented mesoscopy approach is applicable to any target protein that can be labelled for light microscopy, and tissue samples whose dimensions are similar to those used here. We recognise the limitation of using a GFP-tagged GLUT4 transgene in interpreting biological significance to any observed changes, as the transgenic GLUT4 construct is not likely to be transcriptionally regulated in the same fashion as endogenous proteins. Nevertheless, our data provide a useful example of the variety of information that can be extracted from this Mesolens-acquired reconstructed 3D volumes.

The capability of examining defined cardiac regions revealed a significant decrease in GLUT4-GFP levels in the LV of female mice fed a HFD when compared to Control littermates. These data are intriguing considering that females with diabetes have a 50% higher risk of developing fatal coronary heart disease compared to males with diabetes (Huxley et al., 2006). Similarly, studies have shown that patients with HF with preserved ejection fraction are predominantly females, and that females are more likely to have multiple comorbidities, including diabetes and obesity (Cheng et al., 2014; McHugh et al., 2019). However, while animal models are widely used to study the physiopathology of human diseases, the distinctions of diet style-induced cardiovascular disease between males and females remain largely unexplored. Our data highlight the importance of sex equality in research and later the development of adapted treatment.

Ware et al. (2011) and Maria et al. (2015) previously revealed cardiac regional heterogeneity in GLUT4 content under pathological conditions (Ware et al., 2011; Maria et al., 2015). However, from the use of crude cellular samples, the question remained whether variations in protein content represented a global change, or whether distinct architectural features within different regions of the heart may adapt differently. Here we show a skewed distribution of GLUT4-GFP levels along ventricular and septal wall-width within 3D myocardial volumes, which may well underscore changes in contractile activities. Such changes cannot be detected using other currently available approaches. In addition, within the context of the heart, some further notable points revealed by this study include a capacity to visualise and quantify structures that have been classically difficult to study, such as the septum and papillary muscles, and an ability to systematically quantify protein distribution across myocardial walls on the micron-scale in intact tissues.

We introduced a set of quantitative analysis options allowing for better understanding of protein distribution within 3D specimens of intact hearts using ImageJ image processing software (Schindelin et al., 2012). This analysis pipeline remains widely adaptable to diverse molecular targets of interest and the understanding of distinct dynamic patterns within complex tissues. It should be further noted that the 3D datasets captured using the Mesolens provide considerably more information than has been analysed in the current study. For example, the possibility of defining changes in protein expression levels following specific cardiac events such myocardial ischemia in relation to the distinct location of the infract, as well as in tissue adjacent and distal to the injured area could provide real value in understanding whether damage may be reversible or not (Zimmermann et al., 2017). Similarly, quantification of protein expression patterns across endo-, mid- and epi-cardial layers of each chamber of the heart is a possibility that could provide detailed information associated to contractile parameters such as global longitudinal strain (Currie et al., 2005).

Overall, we believe that this novel 3D imaging approach could offer new insight into metabolic changes across the intact heart in models of cardiovascular disease, reconciling research of cardiac remodelling both at a structural and biomolecular level.

## METHODS

### Mice

HA-GLUT4-GFP transgenic mice were obtained from German Diabetes Foundation, Düsseldorf, Germany. These animals (RRID: IMSR_JAX:027496) express GLUT4 containing a hemagglutinin (HA)-tag within the exofacial loop, and a green fluorescent protein (GFP)-tag fused to its carboxyl terminus, under the control of the muscle creatinine kinase promoter. HA-GLUT4-GFP trafficking translocation to the plasma membrane of cardiomyocytes isolated from these mice has been reported and studied (Fazakerley et al., 2009; Lizunov et al., 2012). All animal work was performed under the auspices of Home Office Project Licence (UK) PP5059943 and establishment licence X56B4FB08.

### Diet and Feeding

A total of 14 mice, including 7 males and 7 females were fed with standard chow diet (pelleted; 10% kcal% fat; 824050, DMB Scotland Ltd) up to 5 weeks of age. At 5 weeks, mice were randomised to either Control or High-Fat Diet (HFD) groups. Thereafter, mice in the Control group were kept on a standard diet, while HFD mice were switched to a diet with higher fat content (not pelleted; 60% kcal% fat; 824054, DMB Scotland Ltd) up to the end of the study (20 weeks). Mice were provided with unrestricted access to food and water. Food was only removed for 12h (9am-9pm), pre-blood glucose measurements on weeks 14, 16, and 18. Mice were housed at constant temperature (21±2°C) and humidity of 45-65%, with a 12∶12 hours light:dark cycle.

### Weighting and Blood Glucose Measurement

Following the start of the experiment (week 1), mice were weighed every week (Fig. S2A,B). Blood glucose measurements were first performed 4 weeks after switching diets and repeated every 2 weeks. Blood glucose measurements performed on weeks 14, 16, and 18, were performed following an overnight fast (9pm-9am). Blood glucose measurements were performed using the Alphatrak2 glucometer (Zoetis; Fig. S2C).

### Mice Sacrifice and Heart Collection

Mice were euthanised using 200 mg ml^-1^ Dolethal, which contains sodium pentobarbital. Hearts with the lungs still attached were removed by cutting the ascending aorta and superior vena cava behind the thymus. All tissues were then submerged in Perfusion Buffer (135 mM NaCl, 5 mM KCl, 330 µM NaH2PO4, 10 mM HEPES, 5 mM Na-Pyruvate, 10 mM, 1 mM MgCl_2_ in ultrapure 18.2 MΩ.cm H2O; pH 7.4; autoclaved before use, supplemented with 5 mM Glucose on day of isolation, and filter-sterilised), where the lungs, thymus, and any remaining connective tissues were carefully excised. Hearts were then gently pressed down using the back side of curved tweezers, removing as much blood as possible. Finally, hearts were setup on an Eppendorf lid within a clean petri-dish, where the aorta was canulated and clamped. Hearts were then slowly perfused with 20 mL Perfusion Buffer, followed by 20 mL of 4% Paraformaldehyde (PFA) Solution. The cannula was then carefully removed, and final heart sections were dissected to an appropriate size for Mesolens imaging (maximum 3 mm thickness; for details see *Mesolens Imaging*) using a sharp-end scissors. For all hearts, 1 mm was measured from the apex before measuring and trimming the 3 mm-thick required section. Specimens were then transferred into a 4% PFA-filled Eppendorf and fixed overnight at 4°C, in the dark, with gentle agitation before optical clearing (see Specimen Clearing).

In this paper, we use the term intact when referring to final isolated heart sections, where the core structure remains untouched and individual cardiac chambers undissected, as described above and conversely to the tissue homogenate samples used for immunoblotting analysis (for details see *Tissue Homogenate Preparation*).

### Specimen Clearing

Following fixation, specimens were optically cleared using the iDISCO method (Renier et al., 2014). Hearts were first washed three times in 1xPBS for 30 minutes at room temperature on a rotator and were subsequently dehydrated through a series of methanol solutions, starting with a 20% methanol and distilled water solution, for 1 hour with no agitation. This was repeated in 40%, 60%, 80%, and 100% methanol solutions. Specimens were then left overnight at room temperature in a final 100% methanol solution, without agitation.

Dichloromethane (DCM) was then introduced by placing the specimens in a 1:2 methanol to DCM mixture for 3 hours, followed by two 15-minute washes in 100% DCM at room temperature on a rotator. Finally, the hearts were transferred in an opaque Eppendorf tube and submerged in dibenzyl ether (DBE) until becoming visually transparent (Fig. 1A), at which point specimens were mounted for imaging with the Mesolens.

### Imaging Parameters and Data Acquisition

Individual specimens were secured in custom-designed mounts for long-term imaging with the Mesolens, as described in Clapperton et al. (2024).

A full report detailing the specification and performance of the Mesolens instrument has been published by McConnell et al. (2016). As such, we provide here only a concise overview of the Mesolens, along with modifications tailored specifically for imaging of clarified sections of murine hearts.

Fig. 6 illustrates a schematic of the multimodal setup employing the Mesolens. An illumination source consisting of a 488 nm laser (Multiline Laserbank, Cairn Research) was utilised for excitation of the GFP fluorescence (HA-GLUT4-GFP) of the specimens. A total laser power of 4 mW at the specimen plane was maintained during imaging.

**Figure 6.**
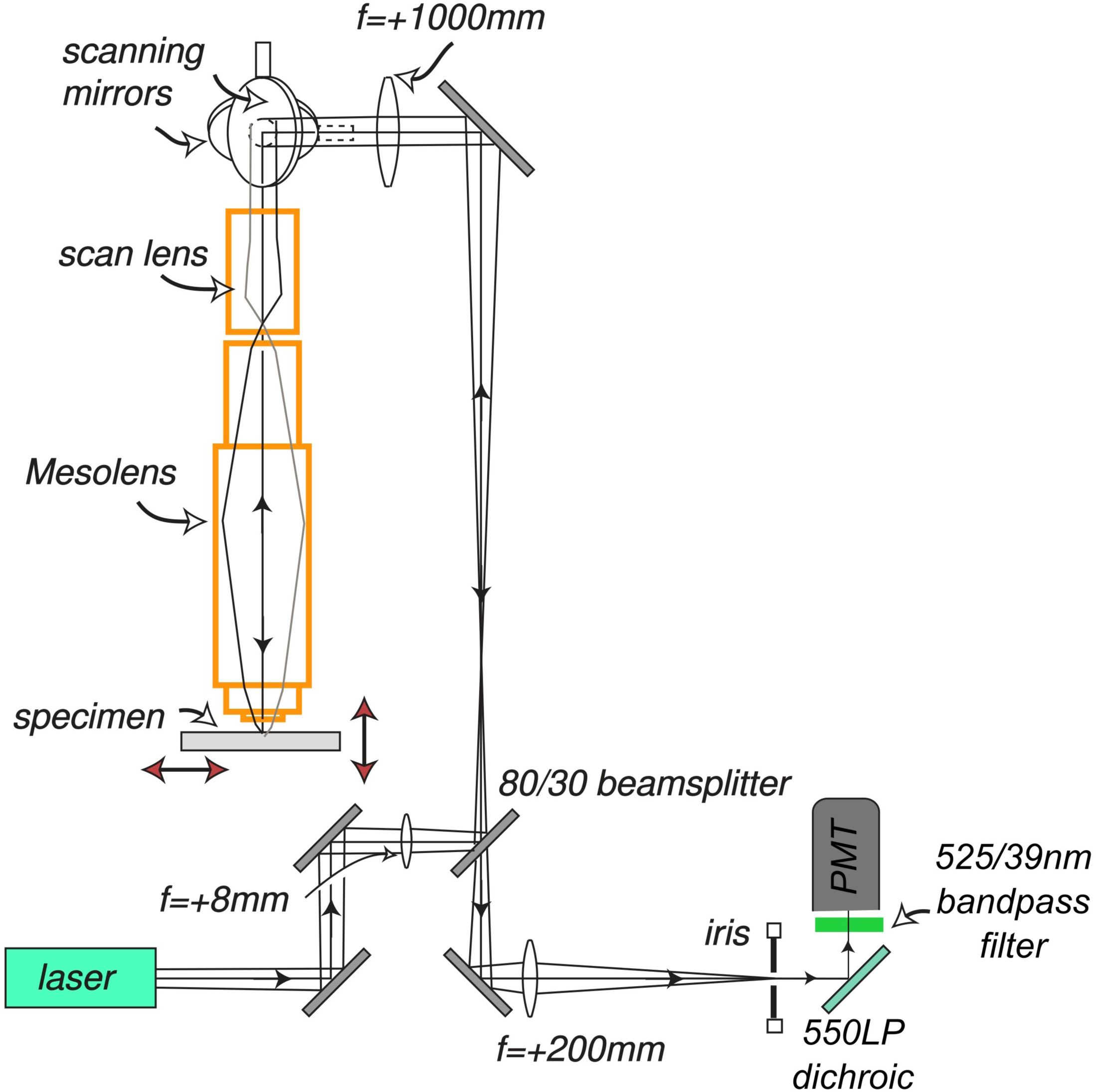
Schematic diagram of point-scanning confocal Mesolens for imaging of GLUT4-GFP in ultrathick sections of cleared mouse heart. A 488 nm laser was used for excitation of fluorescence from the GLUT4-GFP. A single confocal iris was used in the detection path. A single photomultiplier (PMT) was used for detection of the fluorescence contrast together with a 550LP dichroic and a 525/39 nm bandpass filter to minimise backscattered laser light and transmit only fluorescence from GLUT4-GFP.

The Mesolens was used as a point-scanning confocal laser scanning instrument for imaging of all specimens. A 5 mm × 5 mm lateral field of view was scanned with a sampling rate of 1 pixel per micron. While this did not satisfy the Shannon–Nyquist criterion (Shannon, 1949), data proved to provide good optical sectioned images, retaining high spatial resolution without the need for usually larger data volumes to be downsized for faster analysis. The pixel dwell time was set to 0.5 µs, giving an acquisition time of approximately 50 seconds per single optical section of the full 5 mm × 5 mm field. Residual back-scattered 488 nm laser excitation was rejected in the detection path using a dichroic mirror (DMLP505R, Thorlabs, USA) that transmitted wavelengths longer than 505 nm. A bandpass filter (MF525-39, Thorlabs, USA) was placed before the detector to allow only the 525 ± 19.5 nm fluorescence signal to be detected using a photomultiplier tube (PMT) (P30-01, Senstech, UK), for 12-bit detection.

The specimen was moved along the optical axis for imaging at discrete z planes using the confocal method with a computer-controlled *z*-positioning system (Optiscan II, Prior Scientific). To minimise data acquisition time the *z*-step size was set to 5 µm. The system was controlled using an in-house laser scanning software package, ‘Mesoscan’, designed to handle scanned images. Images were stored in the Open Microscopy Environment OME.TIFF format and processed using ImageJ, Excel, and GraphPad Prism software as described below.

### Data Processing and Analysis

The analysis pipeline of image processing, visualisation, and quantification of GLUT4-GFP fluorescent signals described below is summarised in Fig. 2.

Following acquisition, raw Mesolens data were processed using ImageJ image processing software (Schindelin et al., 2012). For each specimen, the individual optical sectioned images were used for reconstruction of 3D heart structures. With a z-step size of 5 µm, average imaging depth was 2.06 ± 0.33 mm (Fig. 1C). 3D volumes were then displayed as 2D images by performing an average intensity axial-projection (Fig. 2A; Image > Stacks > Z Project > Average Intensity).

Heart No.9 (Fig. S2D) was excluded from the analysis process as anatomical structures were visibly damaged.

#### Analysis of Global Regional Fluorescence

The polygon selection tool was then used to manually segment each heart region of interest (ROI), namely the LV, septum, and RV. Each ROI was saved in the ROI Manager (Analyse > Tools > ROI Manager…) and area, as well as fluorescent signals were measured (Fig. 2B). Fluorescent signals, also defined as mean grey values in ImageJ, were calculated from the sum of the grey values of all pixels within the ROI divided by the area of the ROI. Extracted values were then processed using GraphPad Prism 10 (for details see *Statistical Analysis*).

#### Analysis of Transmural Fluorescence

Subsequently, the Plot Profile tool (Analyse > Plot Profile) was used to generate fluorescence intensity profiles across the width (x-axis) of the left ventricular, septal, and right ventricular walls from average intensity axial-projections of Mesolens-acquired 3D reconstructed ventricular volumes (Fig. 2C). In this work, we refer to the fluorescence extracted from these profiles as the transmural fluorescence. For each heart, the line selection tool was used with a linewidth set at 300 pixels and intensity plot ROIs were manually segmented by drawing a line from the LV to the RV, crossing the septum and passing in between LV papillary muscles. Our rationale for using this approach was to mirror one of the most common cross-sectional views of the heart (parasternal short axis) used for assessing cardiac wall parameters and contractile function during *in vivo* ultrasound imaging (Benavides-Vallve et al., 2012; Franchi et al., 2013).

Plot profiles were displayed as 2D line graphs where the x-axis represented the distance along the drawn line and the y-axis was the vertically averaged pixel grey value. In this study, all intensity profiles are displayed as the LV, septum, and RV in this order, from left to right along the x-axis. Transmural fluorescence datasets were extracted as CSV files from ImageJ and subsequently processed in Excel (Version 16.82). Individual left ventricular, septal, and right ventricular transmural fluorescence profiles were isolated from each extracted fluorescence profiles by cutting y-axis datasets at the minimum intensity data point between the LV and the septum, as well as the septum and the RV. The widths of septal and both ventricular walls (Fig. S3B,E) were then extracted from obtained x-axis distances of isolated transmural fluorescence profiles.

Mean transmural fluorescence profiles (Fig. 4A,5A) were obtained by averaging y-axis values of each condition for each heart region, aligning the first y-axis intensity data point of all curves with each other. While misalignments inevitably remain toward the right-hand side of each mean profile due to individual differences in wall width amongst specimens, we remind the reader that mean transmural fluorescence profiles (Fig. 4A,5A) were only used for visualisation purposes and that all statistical analysis were performed on individual isolated transmural profiles (for details see *Statistical Analysis*).

#### Analysis of Lumen Size

Finally, to isolate and quantify the area of the lumen within each heart, average intensity axial-projected datasets were subjected to thresholding (Fig. 2D). Images were first colour-inverted using the Invert LUT lookup table (Image > Lookup Tables > Invert LUT), and threshold segmentations were performed using the Default mode and Red display mode (Image > Adjust > Threshold). Threshold ranges were manually set within each inverted image as such that the lumen edges were fully uncovered. The options Dark Background and Don’t Reset Range were selected, and threshold was applied. Selection of the lumen as a ROI was obtained using the Analyze Particles tool (Analyse > Analyze Particles…) with the following settings: Size (µm^2) = 0-Infinty, Circularity = 0.00-1.00, Show = Masks. The lumen ROI was then selected amongst resulting isolated particles within the ROI Manager, and its area measured. Extracted values were then processed using GraphPad Prism 10 as (for details see *Statistical Analysis*).

#### Visualisation and Presentation

For visualisation and presentation purposes, axial-projected images were colour-coded by fluorescent intensity using the “fire” lookup table (Fig. 2E).

### Tissue Homogenate Preparation

Mice were sacrificed and hearts excised as described above (for details see *Mice Sacrifice and Heart Collection*). Hearts were then dissected to isolate the right and left atria, right and left ventricles, and the septum. Sections were kept on ice, weighed, and homogenised using a T8 basic ULTRA-TURRAX® disperser homogeniser in 100 µL Homogenisation Buffer (250mM Tris-HCl in dH2O; pH 7.4, supplemented with 1x Pierce™ Protease Inhibitor Tablet (ThermoFisher Scientific) and 1:100 Phosphatase Inhibitor Cocktail Set II (Sigma-Aldrich)) per 10 mg of tissue. Samples were then used for SDS-PAGE and immunoblotting analysis.

### SDS-PAGE and Immunoblotting

Protein analysis was carried out using methods and antibodies outlined in (Black et al., 2022). Secondary antibody fluorescence was detected using the LI-COR Odyssey® SA Infrared imaging system. Band intensities on high-quality composite images were then quantified using ImageStudioLite (Version 5.2). Mean pixel grey values of proteins of interest were normalised against GAPDH protein intensities.

### Statistical Analysis

All statistical analyses were performed with GraphPad Prism 10 software. GLUT4-GFP fluorescent signals from individual transmural fluorescence profiles were calculated from the area under curve (AUC). AUC analysis also provided information of peak fluorescence intensity along the x-axis as Peak X. n values and statistical analysis are stated in figure legends. *, **, ***, ****, and ns represent p ≤ 0.05, p ≤ 0.01, p ≤ 0.001, p ≤ 0.0001, and not significant, respectively throughout. All analysis were performed blinded. No statistics were used to determine sample size as we had no evidence base on which to project power calculations.

## ACKNOWLEDGMENTS

We thank all the staff at the Biological Procedures Unit, and Calum Wilson for his advice on statistical analysis.

## DATA AVAILABILITY

Raw datasets supporting the findings of this study are openly available from the University of Strathclyde KnowledgeBase at https://doi.org/10.15129/04700d85-f2b2-40f7-a3a2-58261aabf572.

